# State-dependent changes in perception and coding in the mouse somatosensory cortex

**DOI:** 10.1101/2020.08.24.264127

**Authors:** Conrad CY Lee, Ehsan Kheradpezhouh, Mathew E. Diamond, Ehsan Arabzadeh

## Abstract

An animal’s behavioral state is reflected in the dynamics of cortical population activity and its capacity to process sensory information. To better understand the relationship between behavioral states and information processing, mice are trained to detect varying amplitudes of whisker-deflection under two-photon calcium imaging. Layer 2/3 neurons (n=1436) in the vibrissal primary somatosensory cortex are imaged across different behavioral states, defined based on detection performance (low to high-state) and pupil diameter. The neurometric curve in each behavioral state mirrors the corresponding psychometric performance, with calcium signals predictive of the animal’s choice outcome. High behavioral states are associated with lower network synchrony, extending over shorter cortical distances. The decrease of correlations in variability across neurons in the high state results in enhanced information transmission capacity at the population level. The observed state-dependent changes suggest that the coding regime within the first stage of cortical processing may underlie adaptive routing of relevant information through the sensorimotor system.

**Highlights:**

- Network synchrony and pupil diameter are coupled to changes in behavioral state.
- High behavioral state results in enhanced information transmission capacity at the population level, with neurometric curve in each behavioral state mirroring the corresponding psychometric performance
- Behavioral state and calcium signal in primary somatosensory cortex predict choice outcome.

**eTOC:** *In Brief:* Lee et al. investigates the relationship between behavioral states and information processing in the primary somatosensory cortex. They demonstrate increases in behavioral state results in decrease cortical variability, enhanced information transmission capacity and stimulus encoding at the population level.

## INTRODUCTION

The precision with which sensory neurons represent the environment constrains the quality of subsequent processing in higher cortical areas, ultimately influencing the organism’s behavior. However, the activity of sensory cortical neurons can be fully explained only by considering externally-generated afferent (sensory) signals in conjunction with internally-generated activity in the brain (Erchova et al., 2002; McGinley et al., 2015). This internally generated activity – also referred to as spontaneous activity – depends largely on the behavioral state of the animal. Behavioral state can range from active engagement with the environment to quiet wakefulness, and sleep. Changes in behavioral state are reflected in the population activity of cortical neurons (Sabri and Arabzadeh, 2018). This is often characterized by the level of correlated activity: from desynchronized during active engagement to strongly synchronized during sleep (Harris and Thiele, 2011).

The ecological demands of natural environments vary over time, and animals benefit from tuning neuronal processing to match current behavioral goals (Kayser et al., 2005). How do behavioral states, and the corresponding cortical states, impact sensory coding and perceptual performance? Some studies report an increased sensory response in desynchronized states due to lower noise correlations (Beaman et al., 2017; Engel et al., 2016; Minces et al., 2017; Vinck et al., 2015), whilst others report the opposite (Fanselow and Nicolelis, 1999; Hentschke et al., 2006; Krupa et al., 2004; Sachidhanandam et al., 2013). Here, we investigate how the efficiency of sensory processing and the conversion of sensory information into a decision depend on behavioral state. To achieve this, we trained mice to detect vibrations applied to their whiskers. Rodents are frequently active in darkness and can detect minute vibrations from approaching predators and produce vibration signals to warn other members of the colony (Randall, 2010). To detect small vibrations, rodents can immobilize their array of whiskers to acquire sensory information (Diamond and Arabzadeh, 2013; Diamond et al., 2008a). As head-fixed mice performed the task, we used two-photon calcium imaging to monitor population activity in the vibrissal area of the primary somatosensory cortex (vS1) and thus establish how the dynamics of vS1 populations vary from state to state. We aimed to address the following questions: 1) how does behavioral state affect encoding of sensory inputs by single neurons? 2) How does behavioral state influence cortical population dynamics and synchrony? 3) How do these changes in encoding in turn influence perceptual choice?

## RESULTS

### Detection performance and cortical activity in response to vibration stimuli

Head-fixed mice (n=7) were injected with GCaMP6*f* in the vS1 cortex and trained to perform a whisker vibration detection task (Fig. 1A). A series of pulsatile vibrations was presented via a piezo driven mesh on the left whisker pad at amplitudes of 0, 10, 20, 40, or 80µm. Mice were rewarded for licking the spout on trials with vibration (amplitudes of 10, 20, 40, 80µm); licking in the absence of vibration (amplitude of 0µm) was not rewarded (Fig. 1B). In order to capture time-varying global arousal states, mice were allowed to perform this task for an extended period (median session duration, 52 mins/400 trials; interquartile range: 39-59 mins /300-400 trials). Mice successfully refrained from licking when the vibration was absent (Fig. 1C; the rate of licking on stimulus absent trials was not different from rate of pre-trial licking; *p*=0.136, Wilcoxon rank-sum). In the presence of the vibration, three measures were found to vary in a graded manner with stimulus amplitude. First, mice licked at a higher rate with increasing amplitude (Fig. 1C; ROC analysis - Fig. S1A). Second, they showed faster response times with increasing amplitude (Fig. 1C; Fig. S1B). Third, they showed increased detection rates, with the shape of a compressive sigmoid (Fig 1D). Overall, the behavioral results indicate that the stimulus intensities covered a range from near-threshold to reliably detectable.

**Figure 1.**
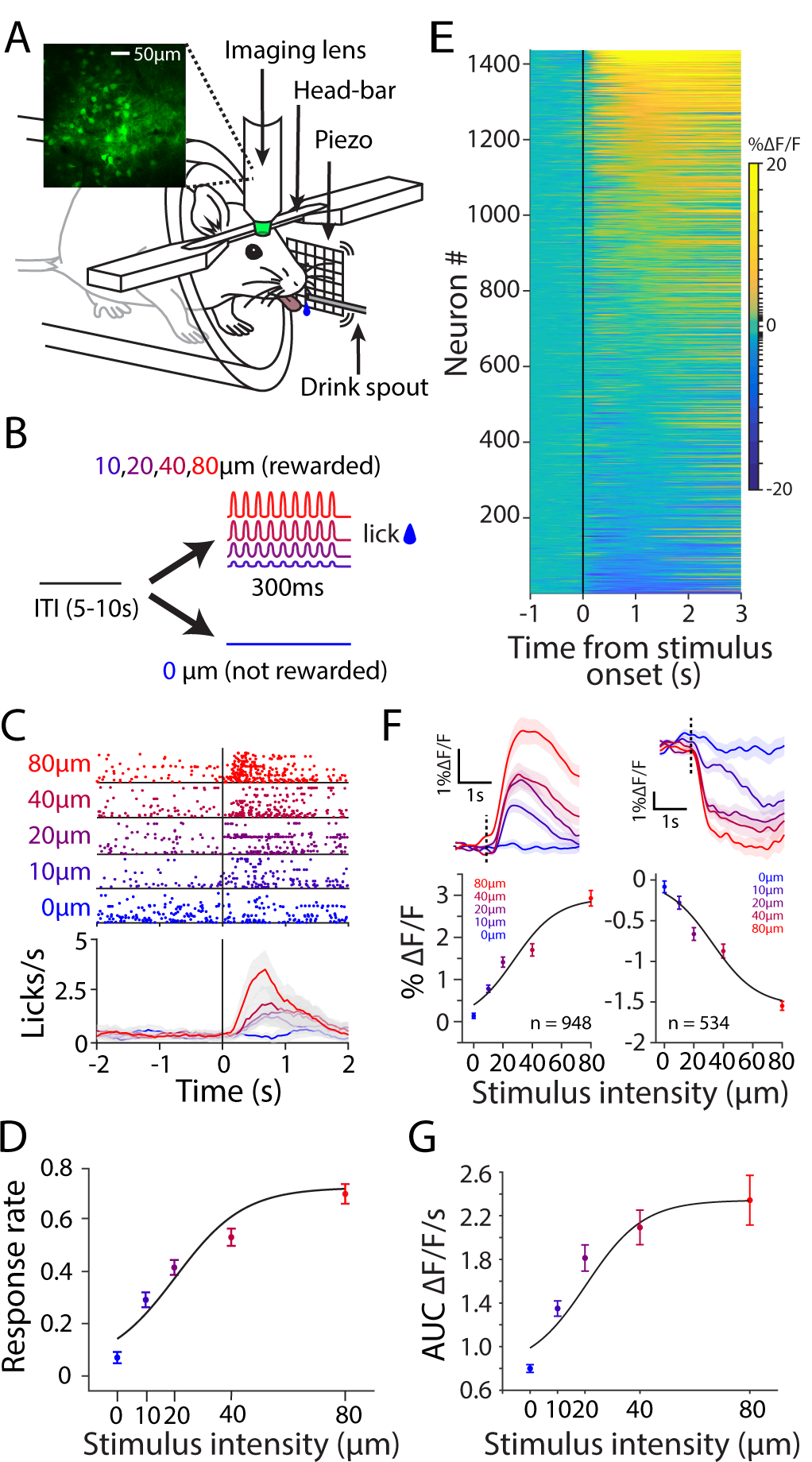
**A**. Schematic diagram of the behavioral setup. As the mouse performed the vibration detection task, we imaged neuronal activity from the vibrissal area of the primary somatosensory cortex using two-photo excitation microscopy (GCaMP6*f*). Stimuli were presented via a mesh place on the left whisker pad and sucrose reward was provided via a capacitive sensing drink spout. **B**. A 300ms 40Hz vibration stimulus was presented at one of five possible intensities (0, 10, 20, 40, 80µm). Licking the drink spout during stimulus presentation (10-80µm) resulted in delivery of sucrose reward. Licking the drink spout at any other time did not result in delivery of sucrose. Each stimulus presentation had an inter-trial/stimulus interval of 5-10s. **C**. Licking profile during an example session. In the raster plot, each lick is represented as a dot and each line represents a trial. Every color represents a different intensity. Average peri-stimulus time histograms are presented with same color notation as in raster plot. Shaded error bars represent SEM. **D**. Hit rate for all mice (n=7) as a function of stimulus intensity. Line represents the best fit of a cumulative Gaussian function. **E**. Every line represents one cell (n=1436). All trials of 80µm vibration are averaged for each cell. Cells are sorted based on their average activity 1 second post-stimulus onset. Scale is logarithmic. **F**. Average calcium trace in response to all stimulus intensities (top), along with corresponding neurometric curve (bottom). Fluorescence values are calculated from 0-1s post-stimulus window. Color notion as in C. Left panels, positive responding cells of E (n=948). Right panels, negative responding cells of E (n=534). **G**. Response modulation as function of stimulus intensity for all cells in E. Responses is calculated by average area under the curve (0-1s post-stimulus window). Continuous line represents the best fit of a cumulative Gaussian function.

We used two-photon calcium imaging to monitor population response of layer 2/3 neurons (example imaging window in Fig. 1A insert). Overall, we recorded a total of 1436 cells across 7 mice (Fig. 1E). Calcium fluorescence was modulated by the onset of the vibration with a heterogeneous profile, including stimulus-evoked increases and decreases in activity. Figure 1E illustrates the heterogeneity by sorting cells based on their average 1s evoked response. Regardless of sign of modulation, across the entire imaged population, neurons showed a graded response to stimulus amplitude: excited neurons (n = 948) became more excited as stimulus intensity increased; inhibited neurons (n = 488) became more inhibited as stimulus intensity increased (Fig.1F). Restricting analysis to significantly responsive cells also showed the same response profile (Fig. S2; excited neurons, n= 343; inhibited neurons, n = 274). To combine both excited and inhibited neurons, we computed the area under the 1-s duration fluorescence trace for all imaged neurons as a measure of stimulus-evoked modulation (Fig. 1G). Area under the curve values exhibited a graded response to the stimulus, in the form of a sigmoidal function with a compressive non-linearity at ∼40µm. The relation of calcium fluorescence to stimulus intensity (Fig. 1G) closely matched the relation of response rate to stimulus intensity (Fig. 1D) (behavioral response function inflection point 11.3 µm; fluorescence response function inflection point 12.1 µm), suggesting fluorescence magnitude as a neurometric correlate of the psychometric detection function.

### Behavioral state affects single-cell coding of stimulus intensity

As mice were allowed to perform the detection task for an extended period each session, we were able to image the same cells over different levels of arousal. Behavioral performance was not static – it waxed and waned throughout each session between periods of high and low detection rates (Fig. 2A, black). On selected sessions, we observed a general slowly progressing decrease in performance. This may reflect changes in the animal’s motivation over time. We examined this by taking into account the false alarm rate (response to stimulus absent trials) across time (Fig. 2A, dash black line). On average, we observed a small but significant correlation between hit rate and false alarm rate over time (r = 0.22, *p*=5.4×10-9 **). This slow time course of motivation could have a different impact on sensory coding than the faster trial-by-trial variability. Overall, across all recorded sessions, mice predominately correctly rejected stimulus-absent trials. Critically, the observed fluctuations in detection rate across time were correlated with pupil diameter (Fig. 2A, orange; see example video – Video. S1). Cross-correlation analysis revealed a moderate coupling between pupil dilation and detection performance (Fig. 2B-right). The temporal relationship was consistently observed across sessions, with pupil diameter lagging behind performance by a median of 9.2 trials (Fig. 2B, left).

**Figure 2.**
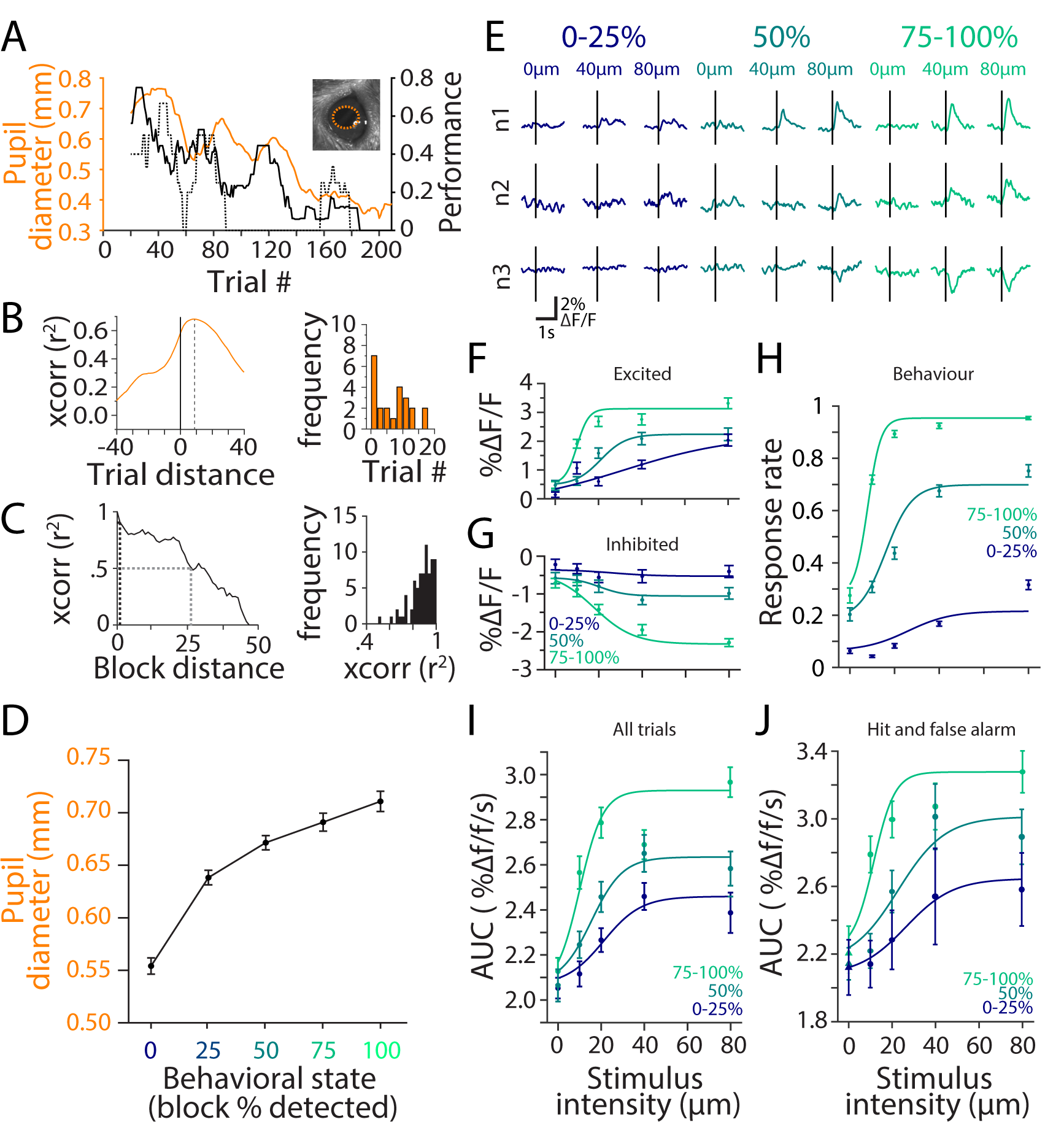
**A**. Changes in pupil diameter (orange), detection performance (black) and false alarm rate (black dashed) of an example session. Lines indicate 10 trial sliding averages of respective measurements. Insert, screenshot of example pupil and estimation of pupil size. **B**. Left, cross-correlogram of performance and pupil diameter in an example session. Right, trial location of peak correlation across all sessions. **C**. Left, auto-correlation of performance in an example session. Black dash line represents correlation coefficient at 1 block distance (5 trials). Gray dash line represents block distance at 0.5 correlation coefficient. Right, frequency distribution of correlation coefficient at 1 block distance across all sessions. **D**. Pupil diameter (average pupil size −5-0s from stimulus onset) as a function of behavioral state. **E**. Average response of 3 example cells to 0, 40 and 80µm at 0-25% (blue), 50% (turquoise), and 75-100% (green) behavioral state. **F**. Neurometric response of top 50% responding cells for 3 behavioral states. Color notation retained from E. **G**. Neurometric response of bottom 50% responding cells for 3 behavioral states. Color notation retained from E. **H**. Psychometric curves averaged across all mice for 3 behavioral states (blue: 0-25%; turquoise: 50%; green: 75-100%). Continuous line represents the best fit of a cumulative Gaussian function to each of the three behavioral states. **I**. Neurometric response of all cells as calculated by area under the curve (0-1s window post-stimulus onset) for 3 behavioral states. Continuous line represents the best fit of a cumulative Gaussian function to each of the three behavioral states. **J**. Neurometric response of all cells as calculated by area under the curve (0-1s window post-stimulus onset) of trials in which animal responded (Hit for stimulus present – filled circle; False-alarm for stimulus absent-filled triangle) in 3 behavioral states. Continuous line represents the best fit of a cumulative Gaussian function to each of the three behavioral states.

Next, we quantified the temporal profile of changes in behavioral performance. Stimuli were distributed into blocks of 5 trials, within which 4 vibration amplitudes (10, 20, 40 and 80µm) were presented in a randomized fashion along with the no-vibration trial (0µm amplitude). This allowed us to quantify behavioral state by calculating the detection rate within each block (0%: no detection; 100%: all four amplitudes detected). Hereafter, we refer to this block detection rate as behavioral state. To capture the temporal dynamics of state changes, we computed the auto-correlation of behavioral state for each session (Fig. 2C, left). Across all sessions, the analysis revealed a high correlation (r = 0.82) between adjacent blocks (5 trials) and an average half width of 27 blocks. Overall, the behavioral state showed a robust correlation with pupil diameter: as the detection rate increased, pupil diameter increased (Fig. 2D) and dilation variance decreased (Fig. S3A). Similarly, hit rate increased as pupil diameter increased (Fig. S3B).

Neuronal activity in vS1 varied in relation to behavioral state. Three example neurons (Fig. 2E) show typical modulations of evoked response with state. Overall, as state transitioned from low (0-25%) to high (75-100%), response magnitude for a given stimulus amplitude increased: excited cells became more excited and inhibited cells became more inhibited (Fig. S4A). How do these response modulations influence the coding efficiency in vS1 cortex? Figures 2F and 2G plot the calcium response functions separately for excited and inhibited neurons. Consistent with the response profile of the example neurons, neurometric functions both for the excited and inhibited populations became steeper as state transitioned from low to high in a manner suggestive of gain modulation. The same profile was found when analysis was restricted to significantly responsive cells (Fig. S4B & C) or when sorting cells based on a smaller, 50ms window (Fig. S4D). Next, we obtained behavioral psychometric curves (Fig. 2H) and compared them with the population neurometric function across all 1436 neurons by calculating the area under the curve values as before (Fig. 2I). The calcium response profiles (Fig. 2I) showed a leftward shift as state transitioned from low to high (inflection points for 0-25%, 50%, 75-100% at 22µm, 15µm, 8µm, respectively). The leftward shift was accompanied by a gain modulation of population neurometric functions: magnitude of change in fluorescence (%ΔF/F) at inflection point increased as state transitioned from low to high (for 0-25%, 50%, 75-100% at 2.28, 2.35, 2.40%ΔF/F, respectively). Again, there was a remarkable correlation between the neurometric and psychometric functions across states (Fig. 2H; inflection points for 0-25%: 26µm; 50%:16µm; 75-100%:8µm). The observation of the elevated neuronal response profile during high-state may have resulted from the higher proportion of hit trials. Previous studies have shown the response of vS1 to be modulated by choice (Poulet and Crochet, 2019; Sachidhanandam et al., 2013; Yamashita and Petersen, 2016; Yang et al., 2016). We performed two additional analyses to better dissociate the sensory component of the evoked response from the motor/decision component. When restricting our analysis to examine only hit trials, we observed a similar response profile (Fig. 2J). In the same light, when restricting our analysis to the first 50ms (to isolate biphasic motor response observed in experiments using electrophysiology (Sachidhanandam et al., 2013; Yamashita and Petersen, 2016), we also observed a similar response profile (Fig. S4E). Finally, given the changes in overall motivation observed by the decreasing false alarm over time (Fig. 2A dash line), we isolated blocks of trials in which false alarm rates were zero (see methods for detail). This analysis also produced similar results (Fig S4F). Overall, as state transitioned from low to high, calcium response profile became steeper in a manner suggestive of gain modulation.

### Behavioral state affects population coding of stimulus intensity

How does behavioral state affect the dynamic interaction between cells and in turn determine the efficiency of the population code? It is well known that stimulus-independent trial-to-trial correlations in activity (also known as noise correlation) limit the quantity of information any neuronal population can carry about the sensory input (Averbeck et al., 2006; Josić et al., 2009; Kohn et al., 2016; Pola et al., 2003). We therefore quantified how noise correlation varied across behavioral states. The analyses revealed a significant drop in pairwise correlation (*p*=2.75×10^−48^**, student t-test) as the state transitioned from low to high (Fig. 3A). This trend was consistent across mice (Fig. 3B, *p*=0.012*, student t-test). Whisker tracking indicated no significant difference in whisker movement between behavioral states (Fig. S5). Noise-correlation strongly varied with cell to cell distance – nearby cells exhibited higher noise-correlation compared to distant pairs (Fig. 3C). From 25µm to 400µm, the drop-off in correlation with distance was steeper and dropped to a lower plateau in the higher behavioral state.

**Figure 3.**
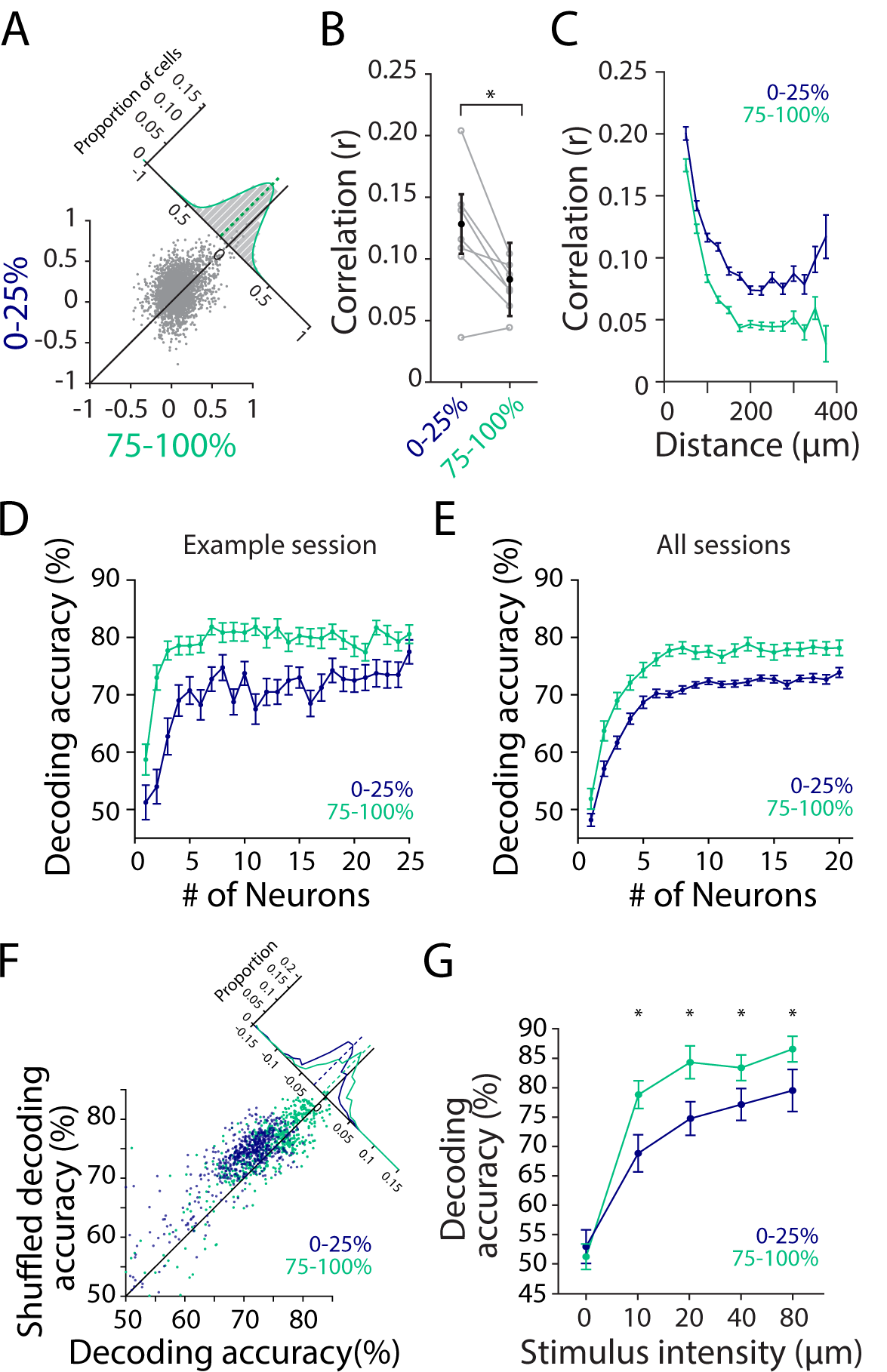
**A**. Noise correlation of cells in example session between 0-25% (blue) and 75-100% (green). Cells in 0-25% blocks exhibit higher noise-correlation than 75-100%. Insert indicates histogram distribution from line of equivalence. **B**. Noise correlation for each mouse between 0-25% and 75-100% (gray). Average noise correlation averaged across all mice between 0-25% and 75-100% (black). **C**. Noise correlation averaged across all mice as a function of cell distance. Blue: 0-25%; Green: 75-100%. Error bars represent SEM. **D**. Linear decoding accuracy of an example session for 20µm between 0-25% (blue) and 75-100% (green) as a function of number of cells. Error bars represent SEM. **E**. Linear decoding accuracy across all sessions for 20µm between 0-25% (blue) and 75-100% (green) as a function of number of cells, up to median population size (20 cells). Error bars represent SEM. **F**. Comparison of linear decoding accuracy of 20µm stimulus between de-correlated trials (by shuffling) and actual data. Each dot denotes the average accuracy of a particular session of a particular population size. **G**. Average linear decoding accuracy for each session across stimulus intensities using all significantly responsive cells recorded simultaneously. Asterisks represents p<0.01, Wilcoxon rank-sum test. Error bars represent SEM.

On theoretical grounds, noise correlation is expected to reduce the efficiency of information transmission by a population (Averbeck et al., 2006; Safaai et al., 2013). We performed linear discriminant analysis to quantify how reliably an ideal observer of the population activity could decode stimulus intensity. Given the levels of noise correlation (Fig. 3A-C), we expect greater information transmission efficiency at higher states. This hypothesis was confirmed by characterizing the accuracy of decoding the presence (versus absence) of the 20µm vibration with growing population size (Fig. 3D - example session; Fig. 3E-average across all sessions). For the same population of neurons, decoding accuracy rose more sharply and plateaued at a higher decoding performance in high state (green) compared to low state (blue). To examine the contribution of noise-correlation to decoding accuracy, we decorrelated the activity of neurons by shuffling trials. As shown in Figure 3F, decorrelating responses led to a significant increase in decoding accuracy for 20µm stimuli in both low-state (blue; *p*=4.31×10^−76^ **, Wilcoxon sign-rank) and high-state (green; *p*=3.04×10^−14^**, Wilcoxon sign-rank). However, the decorrelation-induced increase in accuracy was significantly greater in low-state than in high-state (*p* =4.57×10^−32^ **, Wilcoxon rank-sum). Finally, the enhanced population coding in high state was systematically observed across imaging sessions with various population sizes and was present across all stimulus intensities (Fig. 3G).

### Calcium responses reflect choice outcome

Sensory-evoked activity in vS1 might be expected to predict the subsequent perceptual choice. We asked how the neuronal responses correlate with the mouse’s upcoming behavior, focusing on hit and miss trials. With data from all behavioral states pooled, average response magnitude on hit trials was significantly larger than on miss trials for both excited and inhibited cells (Fig. 4A, 500ms window post stimulus, excited: *p=*1.93×10^−21^*\*\**; inhibited: *p=*3.08×10^−8^,** Wilcoxon rank sum). Next, we asked how neuronal activity can dissociate between hit and miss outcomes in different behavioral states (Fig. 4B & C) (ungrouped categories shown in Fig. S6A). We observed a greater difference between hit and miss response magnitudes in high behavioral states (75-100% detection blocks) compared to low behavioral states (0-25% detection blocks). We quantified the state-related differences in cortical activity after controlling for trial outcome (hit versus miss). Hit trials during high state elicited a significantly greater neuronal response magnitude (both excitation and inhibition) than hit trials during low state (excitation: *p*=1.33×10^−10^**; inhibition: *p*=1.40×10^−10^**). Similarly, miss trials during high state elicited a significantly greater neuronal response magnitude than miss trials during low state (excitation: *p*=2.6×10^−6^; inhibition: *p*=2.9×10^−10^**). From Figure 1D & G, detection rate and calcium response of vS1 neurons is modulated by the strength of the stimulus. Therefore, the proportion of stimulus intensities contributing to hits and miss trials could be different between high and low state. Nevertheless, further analysis examining calcium response across stimulus intensity for hit and miss trials in different behavioral state provided similar results. Overall, hit trials produced larger calcium response than miss trials across stimulus intensities (Fig. 4D). On a similar note, we performed a complementary analysis in which behavioral state was calculated in the absence of the current trial outcome (see Methods for detail). This analysis also produced similar results (Fig. S6B). Finally, we examined neuronal activity of trials in which the stimulus was absent. For both correct rejection and false alarm trials, there was no significant difference in calcium response between high behavioral states (75-100% detection blocks) and low behavioral states (0-25% detection blocks) (False alarm: *p* = 0.6635; correct rejection: *p* = 0.6256, Wilcoxon rank-sum; Fig 4C). Together, these findings imply that the changes in behavioral performance may be defined by the quality of stimulus encoding within vS1.

**Figure 4.**
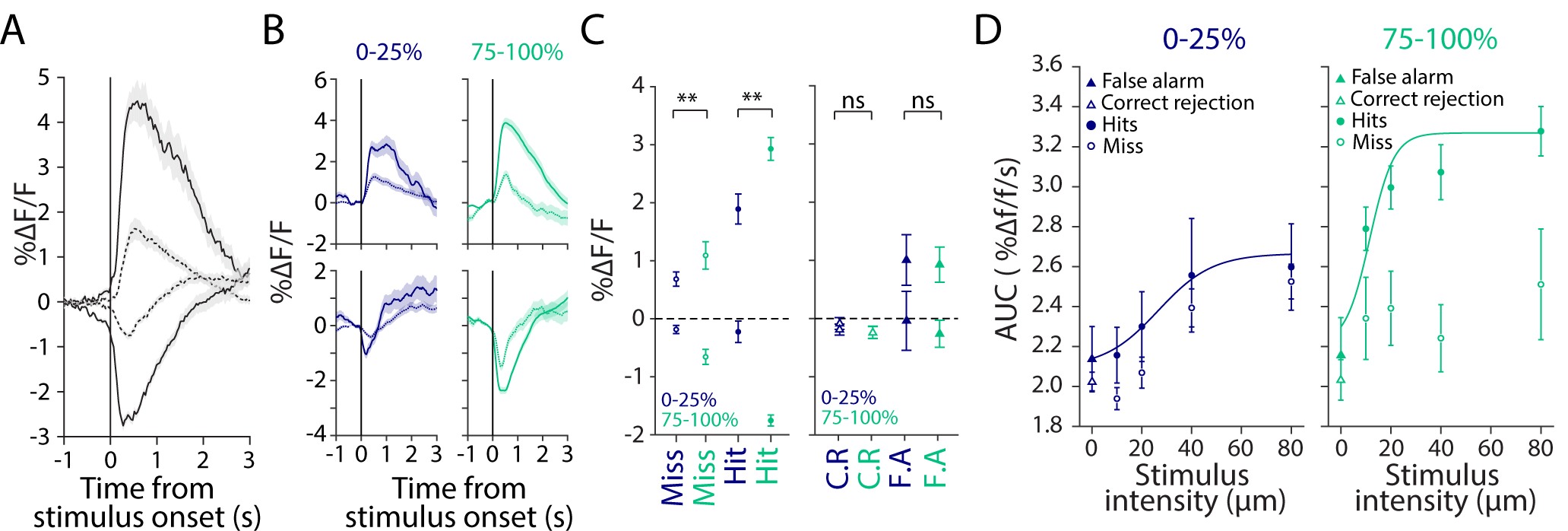
**A**. Average traces of top 25% and bottom 25% responding cells in response to hit (solid line), and miss (dash line) trials. Shaded error represents SEM. **B**. Average traces of top 25% and bottom 25% responding cells to hit and miss trials in low behavioral state (0-25%, blue) and high behavioral state (75-100%, green). Shaded error represents SEM. **C**. Left: 1 second average response of top 25% and bottom 25% responding cells for hit and miss trial in low behavioral state (0-25%, blue) and high behavioral state (75-100%, green). Right: 1 second average response of top 25% and bottom 25% responding cells for correct rejection and false alarm trial in low behavioral state (0-25%, blue) and high behavioral state (75-100%, green). Error bars represents SEM. **D**. Average calcium response profile of cells as calculated by area under the curve (0-1s window post-stimulus onset) for hits (filled circles), miss (open circles), false alarm (filled triangle) and correct rejection (open triangle) across 3 behavioral states.

## DISCUSSION

We investigated the relationship between behavioral state, sensory evoked responses in single neurons, and the dynamics of neuronal population activity in head-fixed mice performing a vibration detection task. Mice reported the presence of a whisker vibration stimulus (0-80µm amplitude) by licking a reward spout and withheld licking during the absence of a vibration stimulus (0µm). In order to capture the transitions between different behavioral states, we allowed mice to perform this task for an extended period each day. Simultaneously, calcium fluorescence in the vibrissae area of the primary somatosensory cortex (vS1) was imaged using a two-photon excitation microscope. As the mice transitioned from low to high behaving states, both psychometric and neurometric curves shifted towards lower stimulus intensities. This enhanced detection sensitivity at the level of single neurons was accompanied by a state-induced reduction in correlated activity across neurons.

Studies investigating cortical state have used whiskers and their central processing pathway as a model sensory system due to the ecological relevance of touch in rodents’ exploration of their environment. Using its whiskers, a rodent can quickly obtain sufficient information to complete complex behavioral tasks, such as discriminating between textures (Diamond et al., 2008b; von Heimendahl et al., 2007; Kuruppath et al., 2014; Zuo et al., 2015), detecting and discriminating vibrations (Fassihi et al., 2014; Lee et al., 2016) and localizing objects (Gordon et al., 2013; O’Connor et al., 2010; Yang et al., 2016). Studies have used whisker movement as a proxy for cortical state (Eggermann et al., 2014; Muñoz et al., 2017; Poulet and Petersen, 2008; Poulet et al., 2012) in the absence of a tactile behavioral task, large amplitude of whisker movement was considered as active state whilst no whisker movement was considered as quiet-quiescence. However, when seeking to acquire signals from a moving object (i.e. a vibration), rodents can actively immobilize their whiskers to optimize sensitivity (Diamond and Arabzadeh, 2013; Lee et al., 2016, 2019). It is therefore imperative to consider this “receptive mode” when investigating cortical state in the whisker system.

Pupil diameter change has historically been hypothesized to correlated with changes in brain state (Hess and Polt, 1960) and our results indicate a strong correlation between pupil size and local detection performance (Fig. 2D). The lag of 9.2 trials suggests that whisker vibration detection accompanies alertness in real time, while the sympathetic control of the pupil follows by about 1 minute (the elapsed time of 9.2 trials). The immediateness of detection performance is one justification of taking this measure as a proxy or identifier for behavioral state.

We observed a heterogeneous response to the vibration, with some cells excited and others inhibited. Critically, we demonstrate that behavioral state modulated evoked response in vS1. As behavioral state transitioned from low to high, the amplitude of evoked response in vS1 increased - excited cells became more excited and inhibited cells became more inhibited (Fig. 2 and Fig. S4A). As behavioral state transitioned from low to high, the amplitude of evoked response in vS1 increased - excited cells became more excited and inhibited cells became more inhibited (Fig. 2 and Fig. S4A). Overall, the high-state decreased synchrony (Fig. 3A-C) while enhancing the evoked responses of vS1 neurons. This finding is different from previous observation in the vS1 cortex where the desynchronized state produced lower evoked responses of cortical neurons (Otazu et al., 2009; Sachidhanandam et al., 2013). The difference may be due to parameters of our behavioral task which demands the whisker system to operate in the receptive mode, actively keeping whiskers stationary (Diamond and Arabzadeh, 2013). The increased amplitude of evoked inhibition in the high behavioral state was also an interesting finding. Fast spiking parvalbumin expressing interneurons receive strong direct sensory input from the thalamus and are responsible for feedforward inhibition (Sermet et al., 2019; Yu et al., 2019). The increased inhibition in high state could be attributed to enhanced activation of these fast spiking interneurons. Alternatively, as excitatory local neurons (e.g. L2/3 and L4 spiny neurons) become more responsive to sensory stimulation, they supply a greater input to inhibitory interneurons, which in turn, may result in stronger inhibition upon their targets. The observed spectrum of excitation and inhibition would therefore depend on the complex interaction between excitatory and inhibitory input a particular cell receives (Baker et al., 2019; Okun and Lampl, 2008; Taub et al., 2013; Zagha et al., 2016). We speculate that the simultaneous and opposite rescaling of the response magnitude of excited and inhibited neurons reflects a mechanism that conserves homeostatic balance across the range of states that sensory cortex naturally cycles through (Xue et al., 2014; Zhou et al., 2014).

The increase in evoked response amplitude was stimulus intensity-dependent while maintaining this inhibition-excitation balance (Fig. 2F & 2G). In the literature, the relationship of state and sensory representation has been unclear. Some studies report an increased sensory response in desynchronized states due to lower noise correlations (Beaman et al., 2017; Engel et al., 2016; Minces et al., 2017; Vinck et al., 2015), whilst others report the opposite (Fanselow and Nicolelis, 1999; Hentschke et al., 2006; Krupa et al., 2004; Sachidhanandam et al., 2013). In our study, evoked responses increased in a non-linear fashion, improving detection sensitivity as animals transitioned from low to high state, mirroring the behavioral performance as mice transitioned from one state to another (Fig. 2H & 2I). Previous research using electrophysiology shows that when a mouse is engaged in a detection task, neurons in vS1 show a biphasic response to the whisker stimulus. This biphasic response is comprised of an early (< 50ms) sensory component and a late (50-300ms) response strongly modulated by the animal’s response (Sachidhanandam et al., 2013; Yamashita and Petersen, 2016). The dynamics of calcium signals restricts our temporal resolution. Nevertheless, when restraining our analysis to the first 50ms, the response profile showed a similar effect (Fig. S4E). Further restricting to our analysis to hold behavioral action constant by examining hit trials only also showed similar effect (Fig. 2J). On selected sessions, we observed a general change in motivation and engagement. As shown in the example session in Figure 2A, at the beginning of a session, mice may perform with high detection rates but at the expense of a high false alarm. This is likely to reflect the high motivation to receive sucrose reward at the beginning of a session. Eventually, false alarm rate decreases while the detection rate is conserved and the mouse performs optimally. This slow change in motivation could have an impact on sensory coding. Nevertheless, by restricting analysis to block of trials in which false alarm rates where zero, we continued to observe evoked responses to increase in a non-linear fashion, improving detection sensitivity as animals transitioned from low to high state (Fig S4F).

As behavioral state transitioned from low to high, the neuronal population showed less synchrony as measured by pairwise noise-correlations (Fig. 3A &B). This is consistent with previous findings in which spontaneous fluctuations in firing rates and intracellular potential show large coordinated fluctuations in cortical population (DeWeese and Zador, 2006; Ferezou et al., 2007; Harris and Thiele, 2011; Luczak et al., 2009; Poulet and Petersen, 2008). These packets of population activity in the synchronized state, interspersed with periods of silence, impose a high level of correlated activity between adjacent neurons (de la Rocha et al., 2015; Scholvinck et al., 2015). In addition to increased synchrony, spatial-temporal dynamics (degree of synchrony across distance) can inform the degree of network dependence (Okun et al., 2015). We found that the strength of correlation decreased with distance (Fig. 3C). This profile of correlation strength with distance supports the idea that nearby neurons share similar excitatory and inhibitory inputs. Overall, our findings are consistent with the distance-dependent decline seen in other studies (Rothschild et al., 2010; Sabri et al., 2016). However, typically these studies examine larger-scale spatial-temporal correlations that may be driven by connectivity between adjacent barrels and barrel-septum connectivity (Sabri et al., 2016). Here, we document state-dependent dynamics on the scale of a single barrel column (∼250µm).

We performed classification analysis as a function of stimulus intensity, the size of the coding population, and behavioral state (Fig. 3E & G). In high behavioral states, where there was a lower-level of correlated activity amongst neurons, decoding performance sharply increased and plateaued at a higher level of accuracy than in low behavioral states. This is consistent with previous studies in which desynchronized state improves the signal-to-noise ratio of the neural code by reducing correlated fluctuations in neural activity, thereby allowing more accurate decisions (Cohen and Maunsell, 2009; Mitchell et al., 2009). In general, correlations among neurons pose constraints on the amount of information encoded in the population and on the decoding. Higher correlation implies redundancy in information. In our study, removal of correlated activity by cross-trial shuffling led to a greater increase in decoding performance in low states (Fig. 3F), indicating that the diminished transmission of information in low states originated both from lower average stimulus-evoked signal per trial (Fig. 2) as well as from high noise correlation. The observed sharp increase in decoding may reflects the tendency of calcium imaging towards more active neurons and therefore, most informative neurons. Alternatively, trained expertise in our vibration detection may have resulted in increased population efficiency to decode stimulus presentation compared to a naïve untrained mouse. The sharp increase may also reflect neural heterogeneity where a small but highly informative subset of neurons sufficiently carries most of the information from the observed population (Ince et al., 2013; Panzeri et al., 2015).

There is a complex interaction between cortical state, the degree of whisker movement, sensory evoked responses at the level of single neurons and the correlation in activity across neuronal populations. Whisker movement affects the membrane potential of vS1 neurons and their evoked response to passive whisker stimulation (Crochet and Petersen, 2006). Beyond single-cell responses, pairwise correlation among vS1 neurons is typically higher in quiet, immobile wakefulness compared to active exploration and whisking activity (Poulet and Petersen, 2008). Whilst behavioral state is often determined by the absence and presence of whisker movement (Poulet and Crochet, 2019), a further distinction can be made within periods of no whisker movement to separate epochs of active engagement in detection from periods of passive resting. In our task, when animals are highly engaged (high-state), whisker movements may be actively suppressed in order to optimize stimulus detection (Kyriakatos et al., 2016). In contrast to the high pairwise correlations observed during low-state, active engagement during high state may have decreased pairwise correlations and in turn improved sensory encoding.

Finally, we examined the correspondence between the vibration-evoked activity in vS1 and perceptual choice of the mouse. We found that state modulated both vS1 activity and behavioral choices. Consistent with recent research, vS1 neurons showed robust choice-related activity (Poulet and Crochet, 2019; Sachidhanandam et al., 2013; Yamashita and Petersen, 2016; Yang et al., 2016) – higher responses were associated with hits; lower responses were associated with misses (Fig. 4A-C). The observed choice related activity was further modulated by behavioral state and this was observed consistently across all stimulus intensities (Fig. 4C & Fig. 4D). However, it is important to note that the observed correlation between vS1 and behavior does not necessarily indicate a causal relation. The extent to which various sensory decision tasks require the direct involvement of sensory cortex remains debated. Whilst some studies show whisker sensory behaviors such as gap crossing (Hutson and Masterton, 1986), roughness discrimination (Guic-Robles et al., 1992), object localization (O’Connor et al., 2010), and vibration discrimination (Miyashita and Feldman, 2013) requires vS1, other studies identify cases where vS1 is not required for active detection of objects and passive detection of air puff stimuli (Hong et al., 2018; Hutson and Masterton, 1986). This disparity likely rests on specific differences in the experimental context such as goal-directed versus habitual reflexive behaviors (Yeomans et al., 2002), appetitive versus aversive conditioning (Guic-Robles et al., 1992; Hutson and Masterton, 1986) or specific stimulus parameters such as stimulus duration and reward schedules (Krupa et al., 2001; Miyashita and Feldman, 2013; O’Connor et al., 2010). The capacity of the brain to generate alternative processing pathways or strategies in response to the loss of function of vS1 must also be considered. Nevertheless, if vS1 is not required for sensory decision making, one alternative explanation for the observed results is that neuronal activity in the somatosensory cortex and behavioral outcomes may both be modulated by state independently. State may affect behavioral outcomes via subcortical processing circuits such as brainstem (Tsunematsu et al., 2020) and thalamic nuclei (Sieveritz et al., 2019) and superior colliculus (Crapse et al., 2018; Wang et al., 2020). The state-modulations in subcortical circuits may then be transmitted to sensory cortex producing choice-related activity in vS1(Yang et al., 2016). In this respect, long-range synchronization between brain regions may underlie functional coupling of areas co-engaged in a given task (Melloni et al., 2007). For example, motor cortex feedback influences sensory processing by modulating network state and the coherence between rat sensorimotor system and hippocampus is enhanced during tactile discrimination (Grion et al., 2016). Future experiments can investigate how long-range synchronization and decision outcomes are affected by specific demands of the paradigm and the animal’s engagement in the task.

The circuit mechanism underlying state modulation is varied, with disinhibition being a key local correlate of response modulation (Chen et al., 2015; Gentet, 2012; Jackson et al., 2016). For example, vasoactive intestinal peptide (VIP) expressing inhibitory neurons in supergranular layers of sensory cortex receive corticocortical inputs from motor areas (Lee et al., 2013) or cholinergic projections from basal forebrain (Zagha and McCormick, 2014). The activation of VIP interneurons in turn inhibit the somatostatin (SOM) expressing inhibitory neurons located in L2/3. Consequently, this disinhibits the inhibition that L2/3 pyramids receive from SOM interneurons and thus trigger the cortical network into a more active desynchronized state. Alternatively, a range of neuromodulatory inputs that arrive into the sensory cortex can influence cortical state (Lee and Dan, 2012). For example, noradrenergic afferents originating from the locus coeruleus contribute to state transition and have been shown to increases neuronal excitability in the somatosensory cortex and improve sensory detection and processing (Sabri and Arabzadeh, 2018; Safaai et al., 2015). In future studies of this detection task, it will be important to study the role of specific subtypes of inhibitory neurons in different cortical layers of vS1 to reveal the neural circuits of sensorimotor transformation from whisker stimulus to goal-directed licking.

## Acknowledgements

The experiments were supported by an Australian Research Council (ARC) Discovery Project (DP170100908), NHMRC project grants (APP1124411), and the ARC Centre of Excellence for Integrative Brain Function (ARC Centre Grant CE140100007).

## Author contributions

C.C.Y.L and E.A. conceived and designed the project. C.C.Y.L and E.K. performed the experiments. C.C.Y.L., M.E.D., and E.A. analysed the data. C.C.Y.L., M.E.D., and E.A. wrote the manuscript. All authors edited the manuscript and approved the final version.

---

**Supplementary Figure 1.**
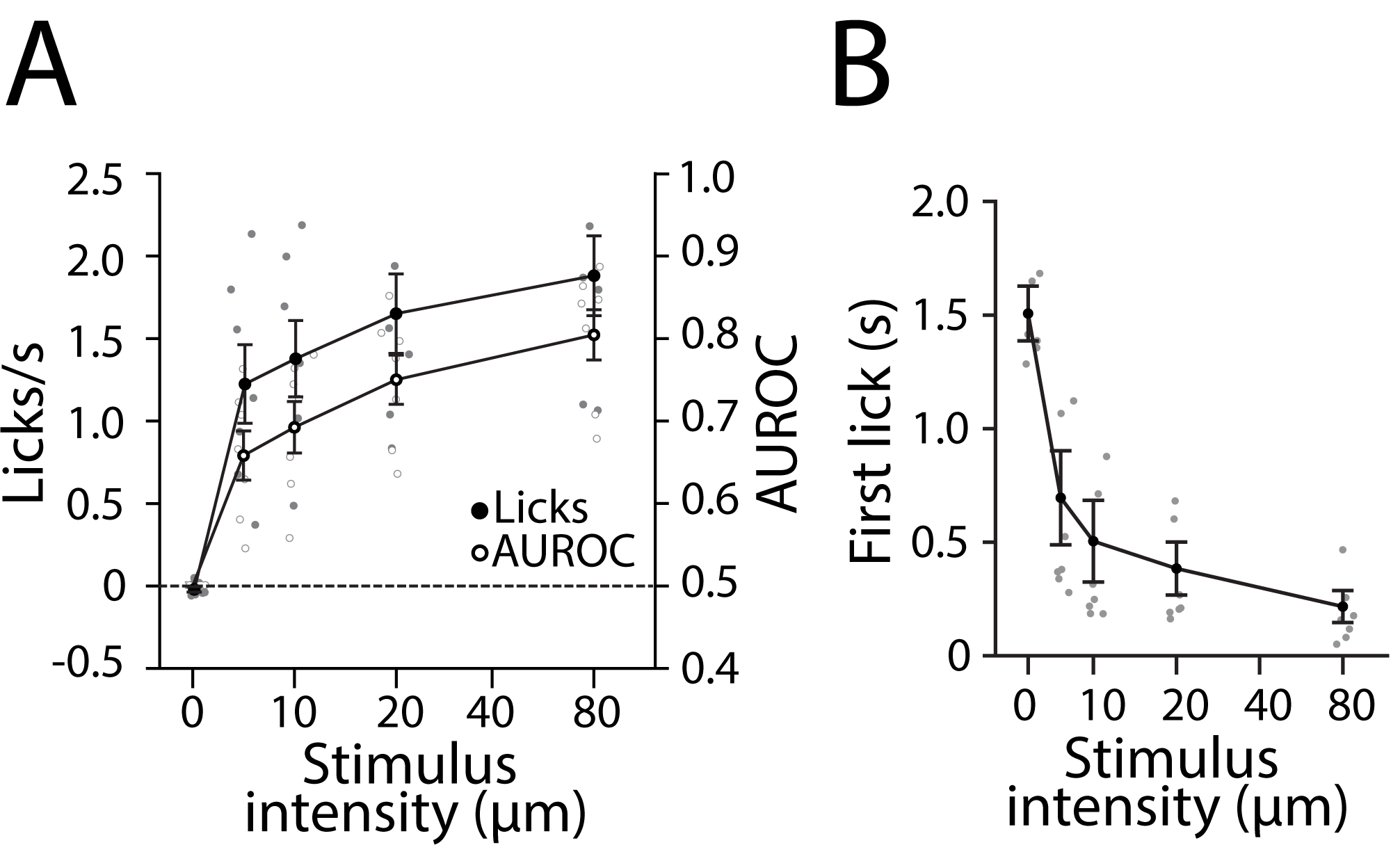
**A**. Lick rate and area under ROC across stimulus intensity. Black open circles represent average AUROC across all mice. Black closed circles represent average lick rate across all mice. Gray open circles represent AUROC for each mouse. Gray closed circles represent lick rate for each mouse. **B**. Reaction time across stimulus intensity. Black dots represent average first lick response time across all mice. Gray dots represent first lick response time for each mouse.

**Supplementary Figure 2.**
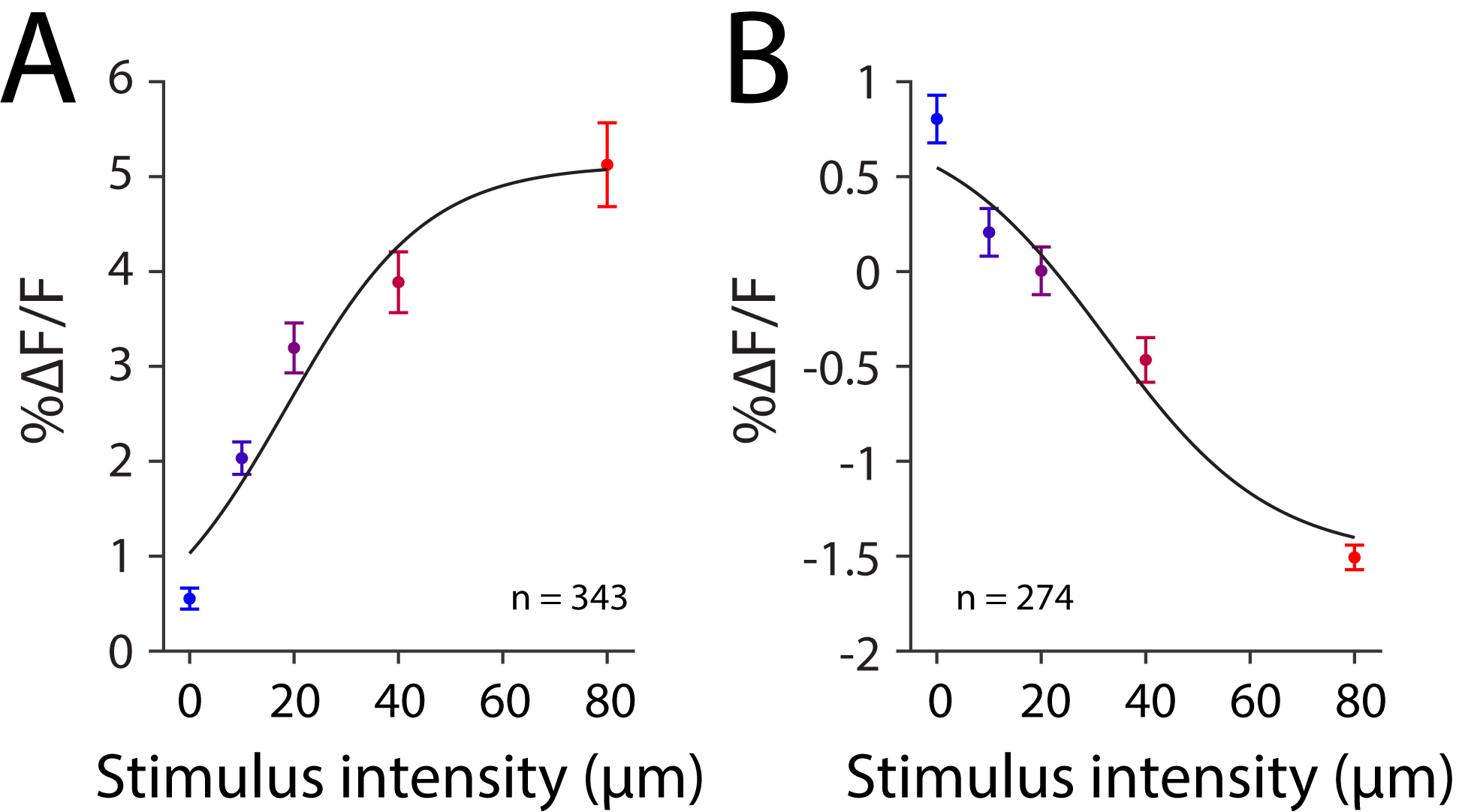
Calcium response functions of significantly responsive cells at 80µm as determined by Wilcoxon sign-rank test. **A**. Average calcium response of significantly excited cells. Line represents the best fit of a cumulative Gaussian function. **B**. Average calcium response of significantly inhibited cells. Line represents the best fit of a cumulative Gaussian function.

**Supplementary Figure 3.**
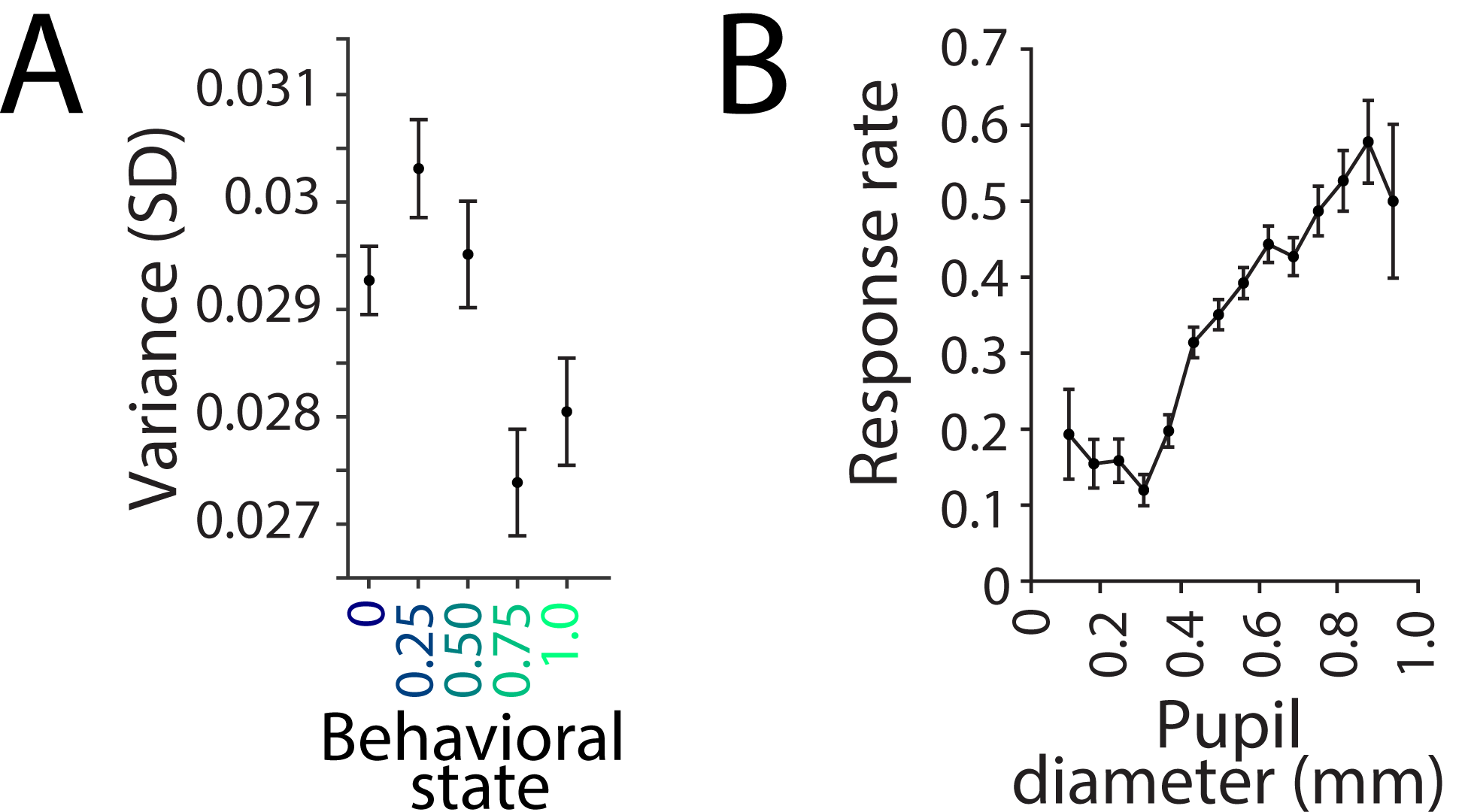
**A**. Variance of pupil diameter as a function of behavioral state performance. Error bars represent SEM. **B**. Hit rate as a function of pupil diameter. Error bars represent SEM.

**Supplementary Figure 4.**
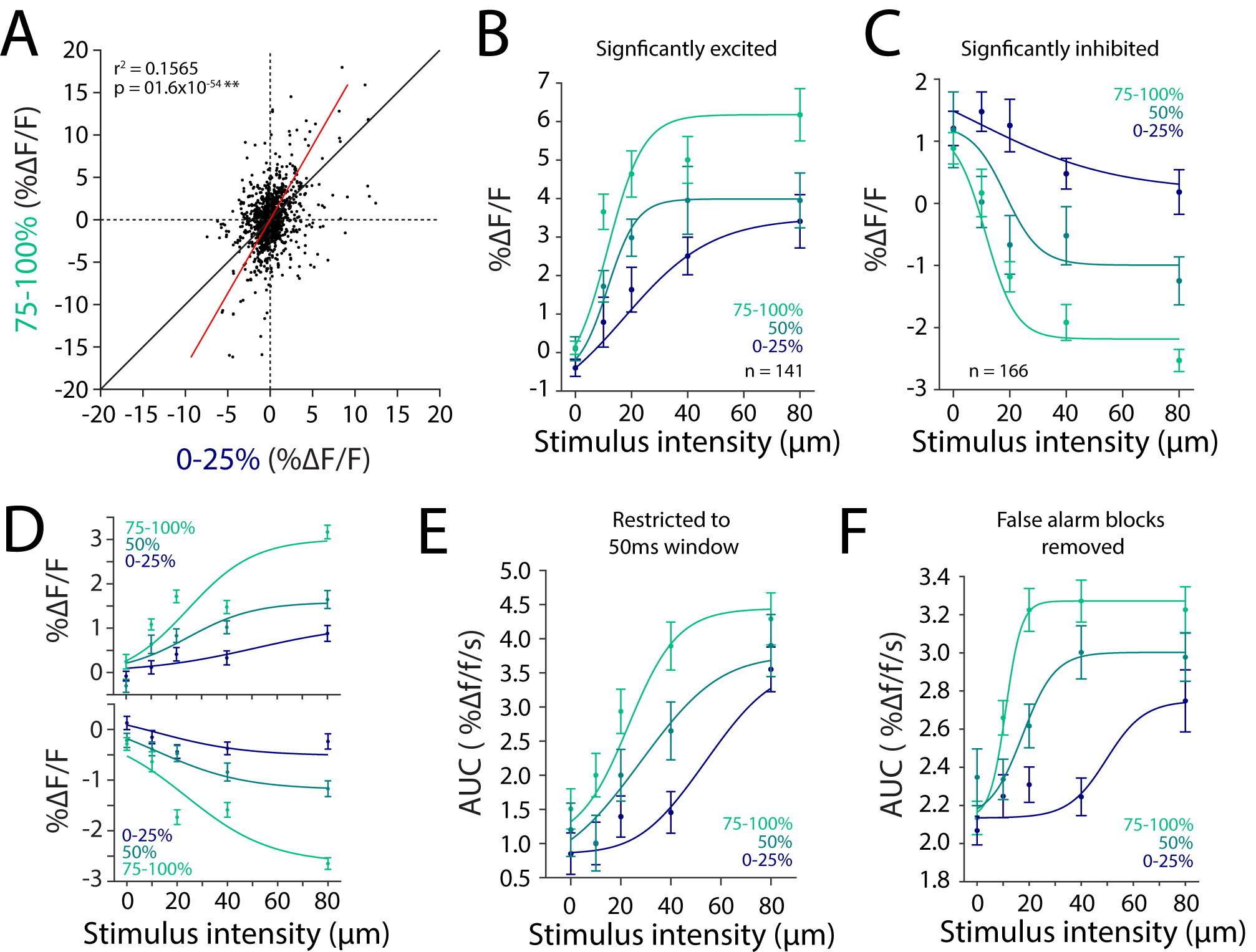
**A**. Scatter plot of evoked response (0-1s post stimulus) across all cells (n=1436) of high state (75-100%) against low state (0-25%). Black line represents line of equivalence. Red line represents linear fit. **B**. Average calcium response profile of significantly excited cells at all 3 behavioral states. Line represents the best fit of a cumulative Gaussian function. **C**. Average calcium response profile of significantly inhibited cells at all 3 behavioral states. Line represents the best fit of a cumulative Gaussian function. **D**. Average evoked response (0-1s post stimulus) of excited and inhibited cells across all 3 behavioral states sorted based on a 50ms window post stimulus. **E**. Average calcium response profile of cells as calculated by area under the curve (0-50ms window post-stimulus onset) for 3 behavioral states. **F**. Average calcium response profile of cells as calculated by area under the curve (0-1s window post-stimulus onset) for 3 behavioral states limited to blocks in which false alarm rate was zero (no response to stimulus absent trials).

**Supplementary Figure 5.**
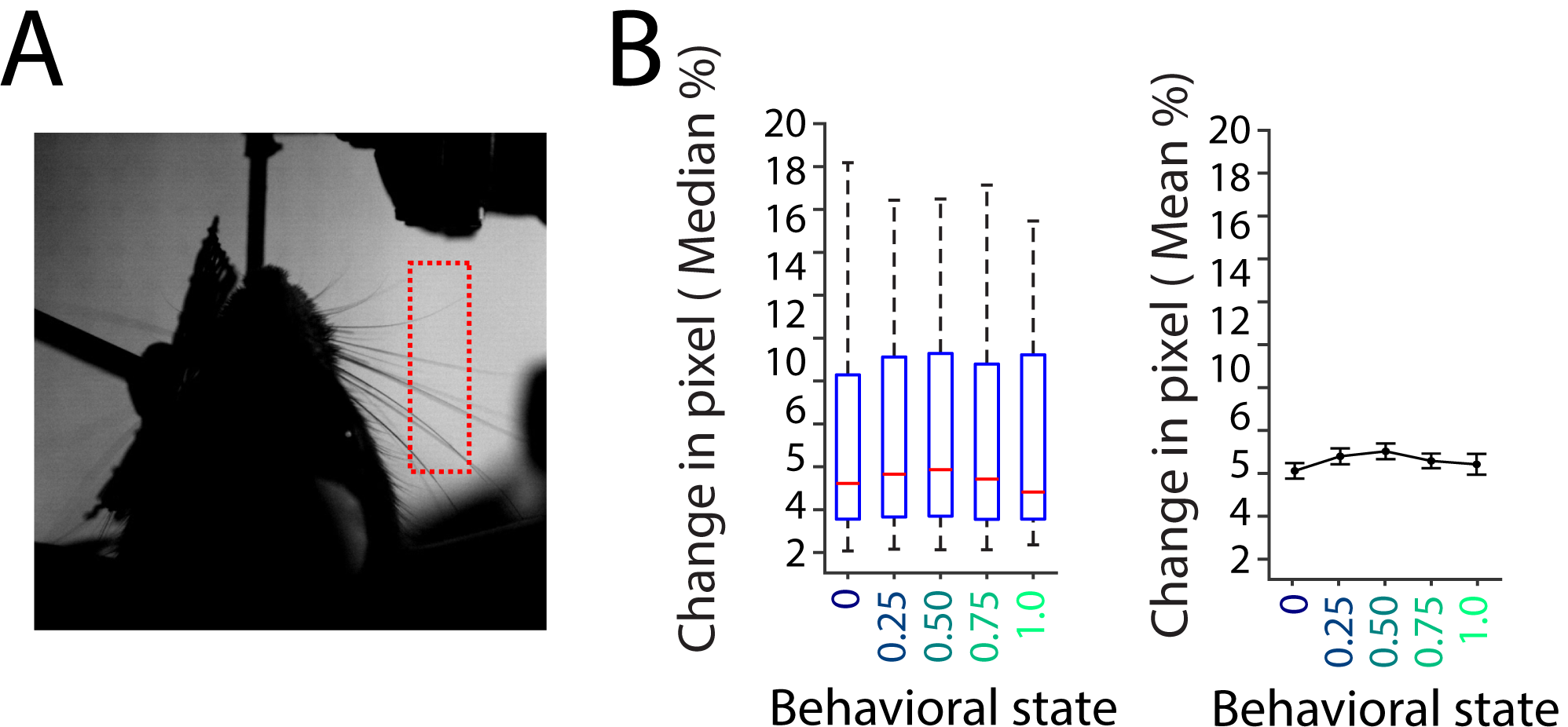
**A**. Screenshot image of whisker tracking video. Dashed area represent region of interest used to calculate percentage change in pixel as a measure of whisker movement. **B**. Left: Whisker and box plot indicating median pixel change (red) in whisker ROI across different behavioral states. Box contains the 25th and 75th percentile; whiskers mark the 5th and 95th percentile. Right: Mean and standard error of mean of pixel change in whisker ROI across different behavioral states.

**Supplementary Figure 6.**
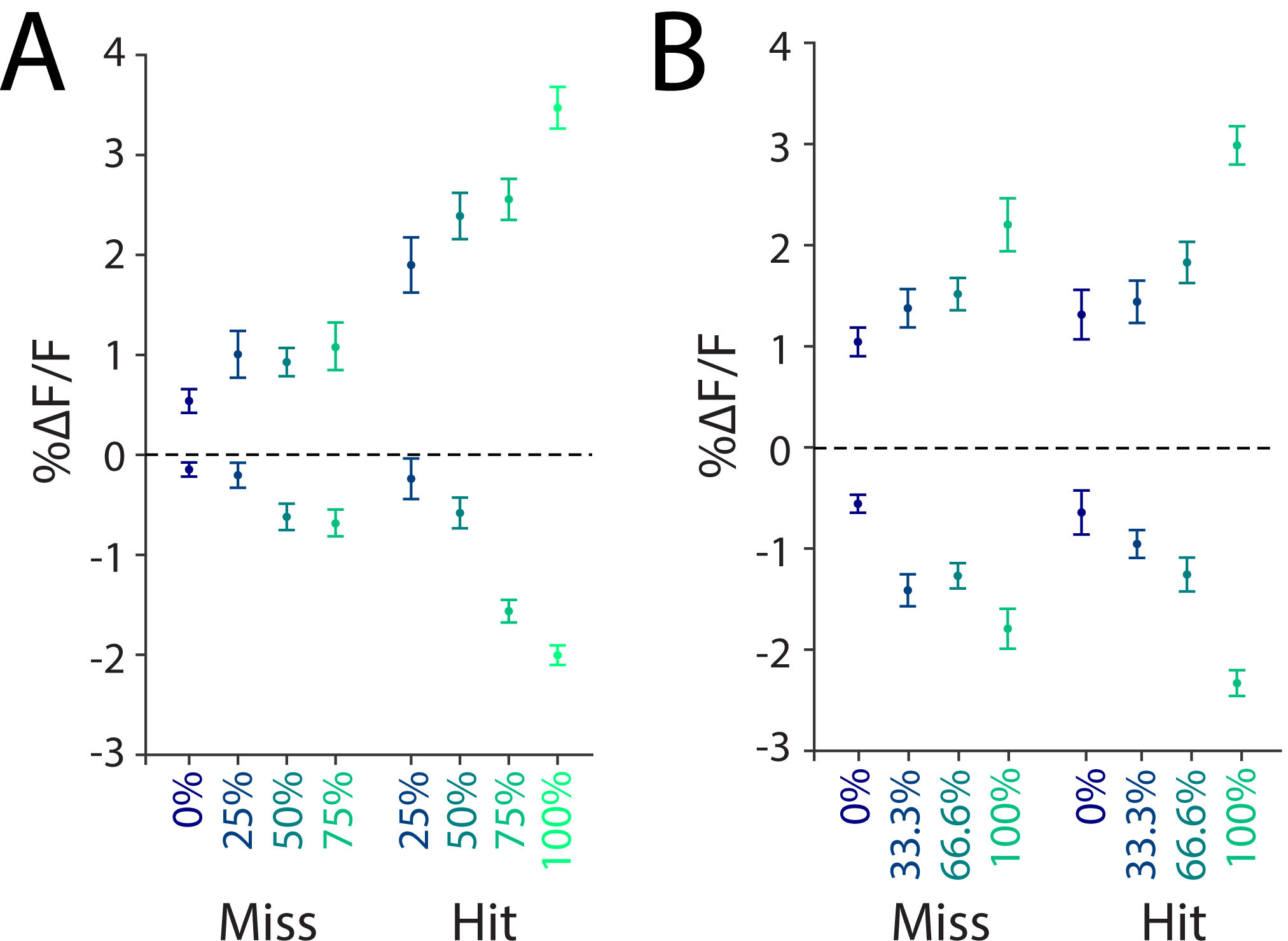
**A**. 1 second average response of top 25% and bottom 25% responding cells for hit and miss trials in all behavioral state categories. Error bars represent SEM. **B**. Same calculations as A for state categories after excluding the response to the current trial of interest (see methods for detail).

## STAR METHODS

### RESOURCE AVAILABILITY

#### Lead Contact

Further information and requests for resources and reagents should be directed to and will be fulfilled by the Lead Contact, Conrad Chun Yin Lee.

#### Materials Availability

This study did not generate new unique reagents.

#### Data and Code Availability

The dataset generated during this study is available at: https://osf.io/de5rh.

The MATLAB codes generated during this study are available from lead contact on request.

### EXPERIMENTAL MODEL AND SUBJECT DETAILS

#### Mice

Subjects were seven 4 week old male C57BL6J with initial weights of 20-25g. All procedures were approved by the Animal Care and Ethics Committee at the Australian National University. Mice were house in independently ventilated and air filtered transparent plastic boxes in a climate controlled colony room on a 12/12 hour light/dark cycle, where lights were turned off at 7pm. Mice were water restricted to motivate animal to perform the detection task. Mice had abundant access to water for 2-3hrs after training sessions and were provided with ad-lib food. All mice gained weight at a normal rate throughout the entire duration of the experiment.

### METHODS

#### Surgery

Mice underwent surgery for viral infection and head-post implantation. They were anesthetized with isoflurane (∼2% by volume in O_2_) and their eyes were covered with a thin layer of Viscotears liquid gel (Alcon, UK) and kept on a thermal blanket to maintain body temperature (Physitemp Instruments). The scalp and periosteum over the dorsal surface of the skull were removed. A circular craniotomy was made over the right barrel cortex (3mm diameter; center relative to Bregma: lateral 3mm; posterior 1.8mm) with the dura left intact. GCamp6*f* (AAV1.Syn.GCamp6f.WPRE.SV40) were injected in 4-6 sites within the craniotomy (4 injections at 32nL per site; depth, 230-250µm; rate ∼92 nLs^-1^) using a glass pipette (15-25µm, tip diameter) via a Nanoject II injector (Drumont scientific, PA). After viral injection, the craniotomy was covered with a glass imaging window 3mm in diameter and 150±20µm in thickness (Warner Instruments, CT). This was glued to the bone surrounding the craniotomy. Custom made head posts were fixed to the skull above lambda using a thin layer of cyanoacrylate adhesive and secured to the skull using dental acrylic. A small well was built surrounding the craniotomy window using dental acrylic to accommodate distilled water required for immersion lens for two-photon microscope imaging.

#### Apparatus

Mice were trained to perform a vibration detection task while head-fixed. All behavioral apparatus was controlled by custom written script in MATLAB (The Mathworks) and interfaced through a data acquisition card (National Instruments, Austin, TX) at a sampling rate of 100kHz. The vibration stimulus was presented to the left whisker pad via an aluminum mesh (2×2cm) attached to a ceramic piezoelectric wafer (Morgan Matroc, Bedford, OH). The mesh was slanted parallel to the animal’s left whisker pad (∼2mm from the surface of the snout). Spacing on the mesh was arranged in a grid with each opening adjacent to one another. Each opening was approximately 150µm apart. At 2mm from the surface of the snout, the diameter of the whiskers of a mouse is approximately 80µm (Carvell and Simons, 2017). Using a microscope, we adjusted the position of the vibrating mesh to reliably engage the maximums number of whiskers to reduce the possibility of poor whisker engagement. Any whiskers that did not enter through an opening on the mesh were rested against the wire structure of the mesh. The vibration stimulus was a series of discrete gaussian deflections. Each deflection lasted for 15ms and was followed by a 10ms pause before the next deflection, yielding a frequency of 40Hz. The total stimulus duration was 300ms. The stimulus amplitude ranged from 0µm, 10µm, 20µm, 40µm, and 80 µm. A custom steel “lick-port” was used to record licks and to deliver 5% sucrose reward and was attached to a micromanipulator to adjust distance for each animal. The lick-port was placed within reach of the tongue at ∼0.5mm below the lower lip and ∼5mm posterior to the tip of the nose. The lick-port was connected to an Arduino Uno acting as a capacitive sensor. The capacitive voltage was sent to data acquisition card and a software threshold was set to determine if a lick was present or absent. The sucrose was delivered via a gravity feed solenoid valve. Mice were placed in an acrylic (4cm inner diameter) tubes such that their heads extended out the front and they could use their front paws to grip the tube edge. A surgically implanted custom head post that extended to the sides of the mice was used to immobilize the animal. Mice were thereby head-fixed in a natural crouching position with their whiskers free to move around the space surround their head.

For recording, the animal was transferred to a two-photon imaging microscope system (ThorLabs, MA) with a Cameleon (Coherent) TiLSapphire laser tuned at 920nm and focused by a water-immersion Nikon objective (x16, 0.58NA). All image acquisition was performed via ThorImage (ThorLabs, MA) and frames were synchronized with the stimulus presentation via the data acquisition card.

#### Training and behavioral task

Training began after surgery and recovery. Animals experienced 4 days of water restriction. During this period, mice were handled to adapt to the experimenter and to the head fixation setup. This involved picking up mice and running the animal through the acrylic tube. With a hemostatic forcep, mice were held in position in the head-fixation post for increasingly longer duration (from 1seconds to 60seconds). Once mice adapted to hand held head-fixation, they were head-fixed into the post for increasingly longer duration (from 1minute to 10 minutes). At each fixation, mice were rewarded with 5% sucrose from a pipette held by the experimenter.

Training began once the animal adapted to head-fixation with one session during which the mouse was rewarded for every lick recorded on the lick-port. When the animal licked, the vibration stimulus was presented for 1 second. By the end of the sessions, mice reliably triggered and consumed the sucrose reward. At the next phase of training, the vibration stimulus was presented indefinitely, until the mice licked the lick-port to trigger release of sucrose. The mice had to trigger the reward three times, after which a 60 seconds no-go period was enforced. During this period, no stimulus was presented and any licks of the lick-port did not release any sucrose. This reinforced the association that licking during the vibration stimulus resulted in a reward, whilst licking when the stimulus was absent resulted in no reward. In three to four sessions, mice reliably triggered reward during the stimulus period and refrained when no stimulus was present. Next, the stimulus period was reduced to 1 second, and the stimulus was either 80µm (go) or 0um (no-go). Between each stimulus presentation, a variable inter-trial interval was enforced, which varied between 5-7 seconds. After mastering this easy version of the task (above ∼85% correct), the stimulus period was reduced from 1 second to 300ms over four sessions and the inter-trial-interval was increased to 5-10 seconds. Once mastered, mice advanced to the final version of the task in which 5 stimulus amplitudes (0µm, 10µm, 20µm, 40µm, 80µm) were introduced. Amplitudes were pseudo randomized in 5 trials blocks, such that each block contained all possible stimulus amplitude. Mice performed the detection task for an extended period each day (∼400 trials), in order to obtain different periods of global arousal states. This was critical as mice experienced the same stimulus intensities and have the same opportunity to response across the entire session.

#### Whisker Tracking

A high speed camera (Mikrotron EoSens CL, Unterschleissheim Germany) was placed above the whisker pad capturing 1000 frames per second at a resolution of 400×480 pixels. A panel of infra-red LEDs illuminated the floor below the whisker pad. 1 second videos clips were captured centered at each stimulus onset. We quantified whisker movement by first filtering each frame via edge detection and measuring the percentage change in pixel intensity from one frame to another in a 200×50 pixel ROI that contained only the ipsilateral whisker pad (Fig.S5A and Video S2). Whisker movement was quantified from a 300ms window before the stimulus onset. All whisker tracking and behavior was performed in darkness. In total, we captured whisking data for 3 mice over 12 sessions each. On average each session was ∼90mins long, totaling 54hrs of high-speed video data. Pixel change analysis captures both whisking amplitude and velocity. We cannot separate these two factors from our measurement. However, from our observation in Supplementary Video 2, pixel change analysis in Supplementary Figure 5 and tracking via WHISK (Clack et al., 2012), we observed no whisking activity nor differences in pixel change/whisker movement.

#### Pupillography

A camera (DMK22BUC03, The Image Source, Taiwan) capturing at 30 frames per second, was place at an oblique angle focusing on the mouse’s left eye. The infra-red light used to illuminate the floor for whisker tracking was also used to illuminate the pupil. All pupil tracking and behavior was performed in darkness. We took precaution in turning off computer monitors and another other light source that could influence pupil dilation. Pupil area was isolated using a custom written MATLAB code that utilized a combination of frame scaling and circle Hough transformation.

### QUANTIFICATION AND STATISTICAL ANALYSIS

#### Behavioral analysis

Hit trials were defined as the presence of at least one lick 0-300ms post stimulus onset and no licks 300ms before stimulus onset. Miss trials were defined as the absence of lick 0-300ms post stimulus onset. Behavioral stimulus detectability was computed from distributions of lick counts occurring 1s before and after each stimulus onset. A criterion shifted in steps of one lick across the two distributions was used to determine the hit and false alarms of a stimulus condition, thus forming a receiver operating characteristic (ROC) curve. Detectability was expressed as the area under the ROC. State was classified into three categories based on the mice performance by calculating the detection rate within each block (0%: no detection; 100%: all four amplitudes detected): low-state (0-25%), intermediate-state (50%) and high-state (75-100%). Since the 5 stimulus intensities were randomized in blocks of 5 trials, state performance was calculated as the average detection performance for each 5 trial block. The ranges of state performance were then separated into the three categories – 0-25%, 50%, 75-100%.

On selected sessions, we observed a general decline in motivation over time. This was characterized by changes in false alarm (behavioral response to zero stimulus intensity) over time. To account for changes in motivation and task engagement, we restricted our analysis to trials in which false alarm was zero. For majority of trials, this was the case – mice correctly rejected zero stimulus intensity by withholding licking. In each 5 trial block (containing all stimulus intensities: 0, 10, 20, 40, 80µm), we excluded blocks of trials in which the mice responded to the 0um stimulus (false alarm). Calcium response profile for each behavioral state was then calculated with the remaining blocks of trials (Fig. S4F).

As state was defined by stimulus present trials including the trial of interest, the observed effect of outcome modulation may be confounded by the current trial of interest. To mitigate this, we performed an additional analysis to supplement our choice outcome analysis (Fig. S6B). This analysis was performed at each 5 trial block to contain all stimulus intensity: 0, 10, 20, 40, 80µm. Like before, for every trial within each block, we defined behavioral state by calculating the detection performance for all stimulus present trials (10, 20 40 80µm). However, in this complementary analysis, the current trial was excluded in the calculation. This resulted in 4 behavioral state categories: 0%, 33.3%, 66.6%, and 100%.

#### Neuronal response analysis

Image stacks were corrected for motion and regions of interest (ROIs) were selected for each cell in each session using Suite2P (Pachitariu et al., 2016). Raw fluorescence *F* was obtained for each cell by averaging across pixels within each ROI. Baseline fluorescence *F*_0_ was calculated by determining the average *F* in the preceding 500ms time window from stimulus onset. The change in fluorescence relative to baseline, Δ*F*/*F*, was computed by taking the difference between *F* and *F*_0_ and dividing by *F*_0_. For all fluorescence heat maps and average traces, plots were generated by a sliding window average of 10 frames at steps of 1 frame.

#### Cross-correlation analysis

Cross correlation between pupil diameter and performance was calculated from a 5 trial sliding window. Pupil diameter was calculated as the average diameter in each window. To perform cross correlation, both measurements were normalized to vary between 0-1. Correlations were also bias corrected for different lag lengths. To calculate correlation coefficient between neuron pairs, we computed the normalized cross correlogram of each cell pairs during periods of spontaneous activity (0-4s window before stimulus onset). Baseline fluorescence *F*_0_ was calculated by determining the average *F* in the preceding 4-5s time window from stimulus onset. Finally, we calculated the maximum height of the correlogram as a measure of correlation strength.

#### Noise-correlation and classification analysis

For noise correlation, in order to remove any effect of stimulus on correlation, we only performed the pairwise correlations on the spontaneous activity (4s before stimulus onset). For each cell pair, the mean fluorescence activity is calculated on each trial and correlated. Classification analysis employed a linear discrimination classifier, training with 90% of data points and testing on the remaining 10%. The classifier classified calcium response from a 1s window post stimulus onset. This response was baseline corrected (1s pre stimulus onset). We classified stimulus present trials (10-80µm) against stimulus absent trials (0µm). The decoding accuracy for each neuron population size was calculated as the average performance over 100 classifying iterations. Figure 3D shows an example session of decoding 20µm stimulus present trials against 0µm stimulus absent trials. Figure 3E shows the average across all sessions for decoding 20µm stimulus present trials against 0µm stimulus absent trials. To examine the contribution of noise-correlation to decoding accuracy, we decorrelated the activity of neurons by shuffling the trial order within each trial category (ie. shuffling within stimulus present trials and stimulus absent trials for each stimulus amplitude and behavioral state). Trial order was averaged over 100 shuffles for neuron population size iteration. Figure 3F shows the improvement of decoding accuracy for 20µm vs 0µm stimulus across all imaging session and population size. Decorrelating the activity by shuffling trial order improved decoding accuracy in low-state compared to high-state.

